# Antibody Inhibition of Influenza A Virus Assembly and Release

**DOI:** 10.1101/2023.08.08.552198

**Authors:** Yuanyuan He, Zijian Guo, Sofie Subiaur, Ananya Benegal, Michael D. Vahey

## Abstract

Antibodies are frontline defenders against influenza virus infection, providing protection through multiple complementary mechanisms. Although a subset of monoclonal antibodies (mAbs) have been shown to restrict replication at the level of virus assembly and release, it remains unclear how potent and pervasive this mechanism of protection is, due in part to the challenge of separating this effect from other aspects of antibody function. To address this question, we developed imaging-based assays to determine how effectively a broad range of mAbs against the IAV surface proteins can specifically restrict viral egress. We find that classically neutralizing antibodies against hemagglutinin are broadly multifunctional, inhibiting virus assembly and release at concentrations one- to twenty-fold higher than the concentrations at which they inhibit viral entry. These antibodies are also capable of altering the morphological features of shed virions, reducing the proportion of filamentous particles. We find that antibodies against neuraminidase and M2 also restrict viral egress, and that inhibition by anti-neuraminidase mAbs is only partly attributable to a loss in enzymatic activity. In all cases, antigen crosslinking – either on the surface of the infected cell, between the viral and cell membrane, or both - plays a critical role in inhibition, and we are able to distinguish between these modes experimentally and through a structure-based computational model. Together, these results provide a framework for dissecting antibody multifunctionality that could help guide the development of improved therapeutic antibodies or vaccines, and that can be extended to other viral families and antibody isotypes.

## INTRODUCTION

Influenza A viruses (IAVs) are segmented, negative-sense RNA viruses that assemble at the plasma membrane of infected cells (1). The assembly and budding of IAVs involves the coordinated action of the viral surface proteins hemagglutinin (HA), neuraminidase (NA), and the proton channel M2, along with the internal matrix protein M1. HA, NA, and M2 are each abundantly expressed on the surface of IAV-infected cells, and they are packaged into virions during budding with relative stoichiometry of approximately 100:25:1 (2, 3). Although IAV assembly and release is not fully understood, the viral membrane proteins are thought to play differentiated yet coordinated roles in the process. The receptor binding and fusion protein HA forms clusters in the membrane of infected cells and potentially induces membrane curvature (4, 5). NA cleaves the glycosidic linkage between virus particles and infected cells, allowing the release of virions for subsequent rounds of infection (6). M2 contributes to membrane scission and incorporation of the viral genome into budding particles (5, 7). The essential roles that these proteins play during IAV assembly and budding represent vulnerabilities that could be exploited in the development of antiviral countermeasures, including vaccines and therapeutic antibodies. While a number of antibodies have been identified that can function in this capacity (8–10), it remains unclear how broadly conserved this functionality may be.

Antibodies neutralize influenza viruses through multiple mechanisms, including inhibition of viral attachment, blocking of viral fusion in the late endosome, restricting the assembly process, and activation of cell-mediated effector functions (11). While established assays are available to evaluate some of these functions (*e.g.*, hemagglutination inhibition for antibodies that block attachment; microneutralization assays for antibodies that inhibit entry), other aspects of antibody function, including inhibition of virus assembly and release, are more challenging to measure or predict. As a result, antibody discovery and characterization has traditionally emphasized an important but somewhat narrow subset of protective mechanisms. However, recent work demonstrating the potency of non-neutralizing antibodies in the control of infection highlights the extent to which antibodies can function outside the context of direct neutralization (12), raising the possibility that multi-functionality – the ability to restrict virus replication through multiple, complementary mechanisms – may be common. However, quantitative methods that can independently evaluate the distinct contributions that a broad range of antibodies make towards the restriction of virus replication are needed to determine if this is the case.

To begin addressing these questions, we developed a fluorescence-imaging based method to quantify antibody inhibition of IAV assembly and release that is agnostic to both the antibody and the viral protein it targets. Using this method, we observed that a wide range of antibodies targeting different antigenic sites on HA, NA, and M2 are capable of inhibiting virus release. For antibodies targeting HA, we find that inhibition occurs through the crosslinking of antigens - either on the infected cell membrane, or between the viral and cell membrane - in a manner that can be predicted by structure-based models that account for antibody conformational heterogeneity. Inhibition of virus assembly typically occurs at concentrations less than ten-fold higher than the concentrations at which a particular antibody inhibits entry, with some classically neutralizing antibodies that bind to the HA head or the HA stalk inhibiting viral release more effectively than they inhibit viral entry. In addition to reducing the number of viruses released, we find that anti-HA antibodies can also alter the morphology of virions produced during a single replication cycle. Finally, for antibodies that target NA, we find that loss of enzymatic activity accounts for only a portion of their inhibitory effect, and that both the potency and mechanism of these antibodies depends on the HA expressed by the target virus. The framework for understanding antibody function described here may be applied to other viruses that assemble at the plasma membrane of infected cells, and could help guide the development of vaccines that better elicit multifunctional antibodies.

## RESULTS

### An imaging-based assay to quantify antibody inhibition of viral release

To determine the potency of antibodies against IAV surface proteins during virus assembly and release, we developed a fluorescence-imaging based approach to directly count virions released into the cell culture media during a single replication cycle (Figure 1A). Following infection at MOI ∼ 1, we incubate MDCK cells with monoclonal antibodies starting at 2h post-infection (hpi), and we collect viral supernatants at 8 hpi. Released virions are immobilized onto glass-bottom plates coated with *Erythrina Christagalli* lectin (ECL) for fluorescence imaging. This approach is insensitive to high concentrations of HA-specific antibodies (Figure S1A) and gives linear results across a >100-fold range, from 9 PFU/well (the lowest concentration tested) to 1125 PFU/well (Figure 1B). The upper end of this range can be extended arbitrarily by pre-diluting samples for accurate quantification. In comparison to Western blot analysis, particle counting gave a >10-fold lower limit of quantification (Figure S1B). Collectively, these results establish fluorescent particle counting as a quantitative and sensitive assay to measure viral shedding in cell culture supernatants in the presence of a range of neutralizing antibodies.

**Figure 1.**
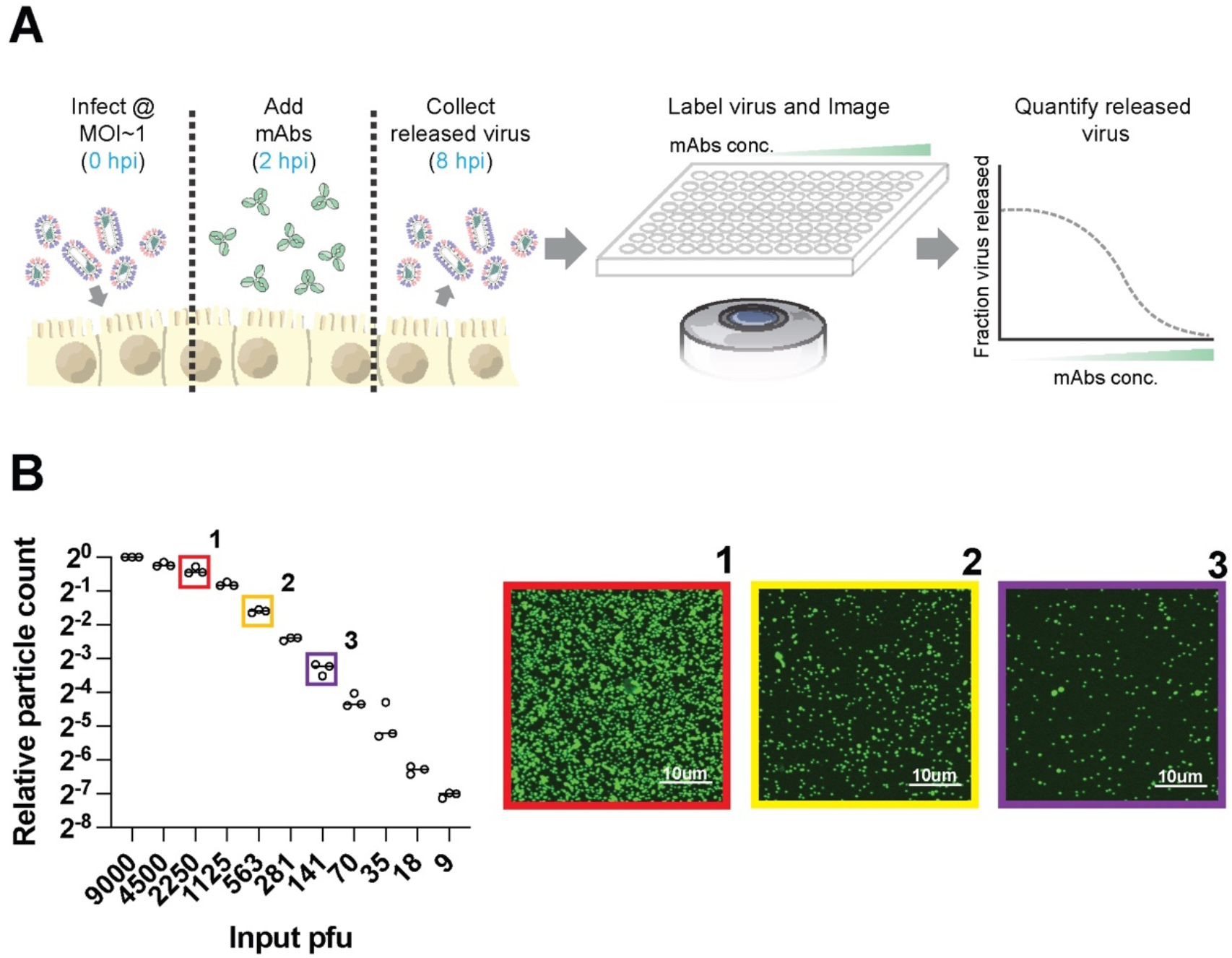
Measuring antibody inhibition of virus release by counting virions. (A) Overview of the image-based assay to measure antibody inhibition of virus release. Cells are infected with influenza viruses at MOI∼1 and incubated with monoclonal antibodies (mAbs) for 2h. Released virions are collected from the supernatant at 8 hpi, labeled, and immobilized for imaging. Segmentation of the resulting images enables quantification of released virions. (B) Sensitivity and linearity of virus particle counting compared to quantification from plaque assays. Individual data points are from three separate serial dilutions of A/WSN/1933 virus starting from 3×10^5^ pfu/ml. Images to the right are from the indicated conditions in *B*. Virus particles are visualized using CR9114 scFv labeled with AF488. Contrast in the sample images is exaggerated to show individual virions.

### Monoclonal antibodies targeting a variety of antigenic sites on HA inhibit viral release

Using this method, we tested the effect of anti-HA antibodies on viral release. HA is the most abundant viral membrane protein on both virions and infected cells, and antibodies targeting a range of sites across the HA surface have been identified and characterized. We selected seven anti-HA antibodies targeting the receptor binding site (RBS) (S139\1 (13–15) and C05 (16, 17)); the central stalk (CR9114 (18), FI6v3 (19), and CR8020 (20)); the trimer interface (FluA-20 (21–23)); and the anchor epitope (FISW84 (24–26)) (Figure 2). These antibodies are broadly reactive, allowing us to compare results against historic strains from different subtypes: A/WSN/1933 (H1N1) and A/HK/1968 (H3N2). We expressed and purified these antibodies as IgG1 isotypes (differing only in their VH and VL domains) and tested their ability to inhibit virion assembly and release. For these experiments, we tested antibody concentrations up to 60 nM, similar to the serum concentration of a dominant clonotype post vaccination (27).

**Figure 2.**
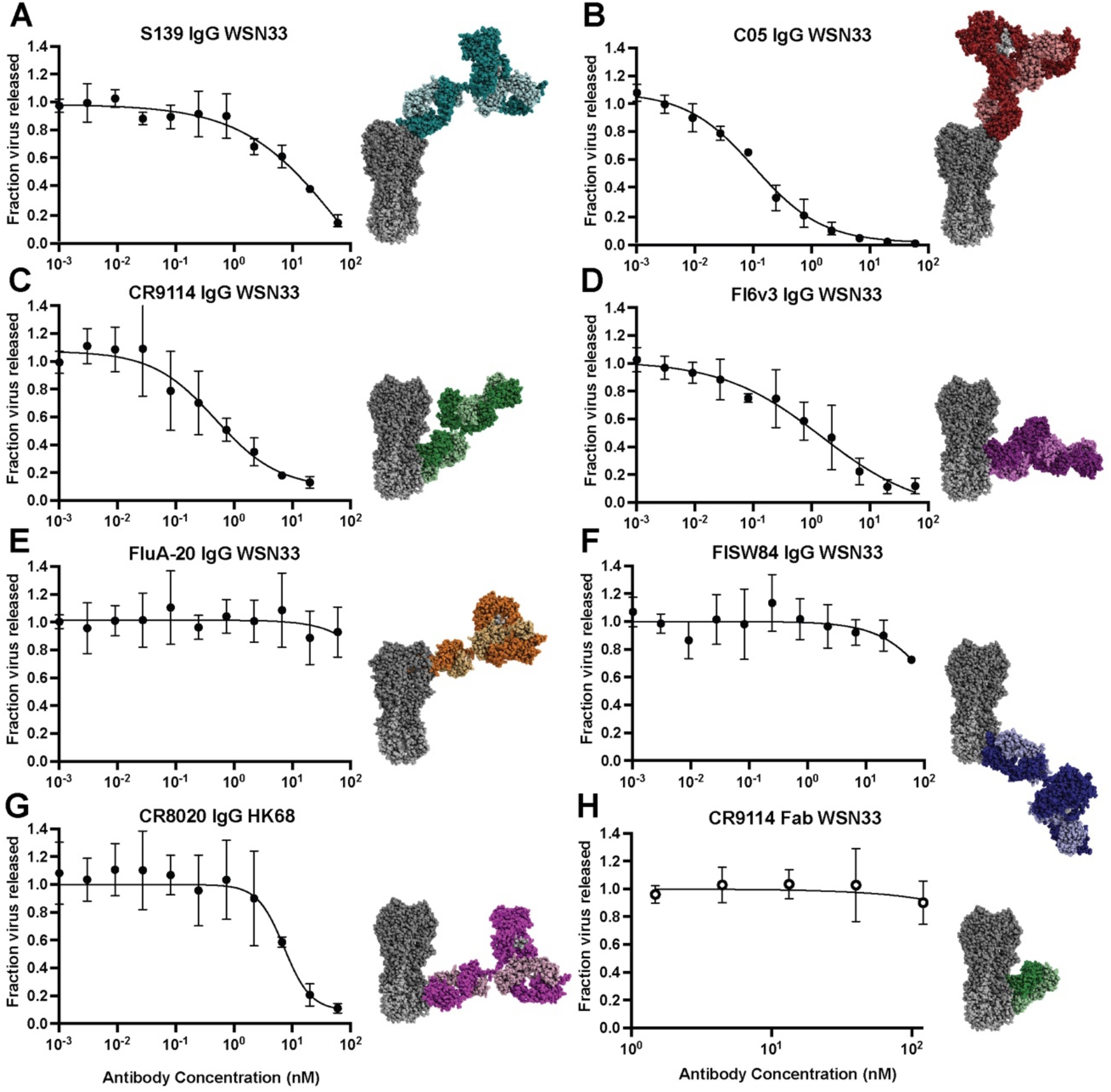
Anti-HA antibodies inhibit influenza virus release from infected cells. (A-H) Neutralization curves showing the fraction of viral particles released from cells infected with A/WSN/1933 (‘WSN33’) or A/Hong Kong/1968 (‘HK68’) as a function of antibody concentration. Each curve is generated from three biological replicates. Error bars show standard deviations and the fit curves are generated by the least squares method using GraphPad prism. Images to the right of each plot show HA (PDB ID 3LZG) in gray with models of full-length IgG1 antibodies bound. Models are obtained by aligning an IgG1 structure (PDB ID 1HZH) to HA:Fab structures (PDB IDs 4GMS, 4FQR, 4FQI, 3ZTJ, 6OC3, 6HJP, and 3SDY). Antibody heavy chains are shown in darker shades with light chains shown in lighter shades.

Antibodies targeting the HA head (S139\1, C05) or stem (CR9114, FI6v3, CR8020) all showed robust inhibition of viral shedding (Figure 2A-G). Inhibitory profiles of these antibodies against filamentous (‘WSN 33 M1Ud’) and non-filamentous (‘WSN33 WT’) strains that differ only in their M segment are largely similar (Figure S2A, Methods). In contrast, these antibodies have greater potency against non-filamentous strains during viral entry (Figure S2B), consistent with previous findings (28). Antibodies that bind to the HA stem have been shown to inhibit NA activity through steric hindrance (29). In testing each HA antibody, we added 0.1 U/ml exogeneous sialidase from Clostridium perfringens (CpNA). Although this treatment is sufficient to restore viral shedding in the presence of the potent NA inhibitor oseltamivir carboxylate (Figure S3B) and has previously been used to rescue viruses completely lacking NA (30), it did not restore viral release in the presence of any of the stem-binding antibodies tested, suggesting that these antibodies are able to restrict viral release through mechanisms other than inhibition of NA.

While FluA-20 IgG and FISW84 IgG both bound to infected cell surfaces (Figure S4A), they did not reach 50% reduction in virion shedding at the maximum concentration tested (Figure 2E&F). We reasoned that this may be due to limited accessibility of the epitopes these antibodies recognize. The FISW84 epitope likely requires tilting of the HA ectodomain to enable binding (24, 26), and the FluA-20 epitope is occluded in the HA trimer, requiring transient opening of the HA head for this antibody to bind. Consistent with epitope accessibility playing a critical role, we observed significant inhibition of viral shedding by FluA-20 against a virus with HA from A/California/04/2009, which readily dissociates into monomers (Figure S4B) (31). The increased potency of FluA-20 against HA from A/California/04/2009 versus HA from A/WSN/1933 - despite the conservation of the five residues with which FluA-20 primarily interacts (21, 22) – highlights the importance of HA trimer stability and epitope accessibility in determining antibody potency in inhibiting virus release.

### Inhibition of viral egress by anti-HA antibodies affect the morphology of released virions

To examine the effect of antibody inhibition on the characteristics of released viruses, we compared particles from the filamentous strain A/Hong Kong/1968 raised in the presence or absence of CR8020 IgG. At antibody concentrations where particle release is decreased by 75%, we observe a 36% apparent decrease in mean HA abundance per particle, measured using a fluorescent Fab fragment from C05 (Figure S5A). This decrease in HA intensity may result from changes in HA abundance per particle, or from interference of CR8020 IgG with C05 Fab attachment. To compare particle size in a way that is independent of labeling intensity, we measured the percentage of viral filaments greater than 1, 2, or 4 μm in length, sizes above the diffraction limit of our optical system (∼300 nm) which can easily be resolved. We find that the percentage of viral filaments above each length threshold successively decreases in the presence of CR8020 IgG relative to the antibody-free condition, suggesting that antibodies can alter both the number and the morphological features of the viruses that are released over the course of infection (Figure S5B).

### Anti-NA antibodies inhibit viral release via mechanisms beyond direct inhibition of enzymatic activity

We next tested two anti-NA antibodies: 1G01, which binds to the active site (10),and CD6 which binds to the interface of adjacent monomers in the NA tetramer (32). Since both antibodies inhibit NA enzymatic activity, we performed these experiments both in the presence and absence of CpNA, which has been shown by us (Figure S3) and by others to compensate for loss of NA enzymatic activity (30, 33). We found that the extent to which inhibition of viral shedding by 1G01 and CD6 IgG could be rescued by CpNA varied depending on the virus’s genetic background. Although all viruses tested express the same NA (from A/California/04/2009), particle release could not be rescued by CpNA in a WSN33 background with mismatched HA (Figure 3A), but was partially restored in a PR8 background with matched or mismatched HA (Figure S6). This suggests that anti-NA antibodies inhibit viral release through mechanisms besides enzymatic inhibition, and that this inhibition varies depending on the genetic context.

**Figure 3.**
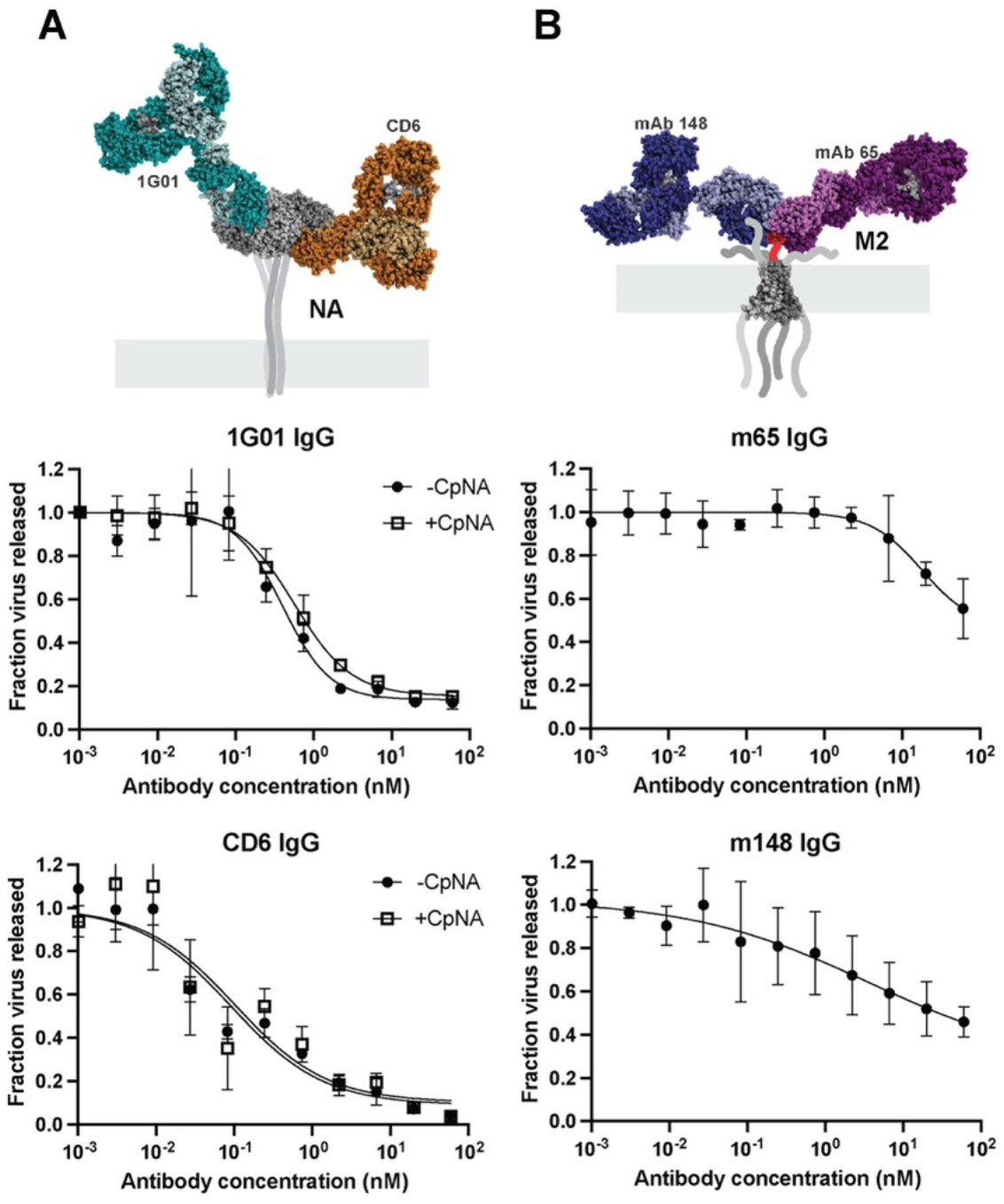
Anti-NA and anti-M2 antibodies inhibit influenza virus release. (A) Neutralization curves for the anti-NA antibodies 1G01 and CD6 IgG against A/WSN/1933 with NA from A/California/04/2009. Experiments are performed with or without 0.1 U/ml exogenous sialidase (‘CpNA’). The model of antibodies bound to NA is obtained from PDB IDs 6Q23 and 4QNP. Curves are generated from three biological replicates. (B) Measurement of anti-M2 antibodies mAb148 and mAb65 against A/WSN/1933. The model of antibodies bound to the M2 ectodomain is obtained from PDB IDs 2L0J, 4N8C and 5DLM. For both *A* and *B*, error bars show standard deviations and fit curves are generated by the least squares method using GraphPad prism.

### Anti-M2 antibodies reduce viral release at high concentrations

Finally, we investigated the ability of two M2-specific IgG antibodies, mAb148 and mAb65, to inhibit virus assembly and release. These antibodies bind to overlapping epitopes in the extracellular domain of M2 (34, 35). While prior work found that an antibody against the M2 ectodomain (14C2) only inhibited the assembly of filamentous strains (9, 36), both anti-M2 antibodies we tested were able to restrict viral shedding of the spherical strain WSN33, but required high concentrations and were generally less potent than HA- and NA-specific antibodies (Figure 3B). Together with our results testing anti-HA and anti-NA antibodies, this establishes inhibition of viral release as a widespread mechanism of protection for antibodies targeting each of the three primary viral surface proteins.

### Crosslinking of HA or NA in cis or in trans contributes to inhibition of viral release

To understand mechanisms that contribute to antibody inhibition of viral release, we compared inhibition profiles of bivalent CR9114 IgG and monovalent CR9114 Fab at concentrations up to ∼100-fold higher than the dissociation constant for both Fab and IgG (∼0.4 nM) (18). In contrast to CR9114 IgG, the monovalent CR9114 Fab showed no inhibition of viral release, confirming that bivalency is important. Together with our finding that exogenous sialidase has limited ability to reverse inhibition of viral egress by anti-NA antibodies but not oseltamivir, this suggests that antigen crosslinking plays a key role in inhibition of viral shedding. This crosslinking could occur between antigens within the same membrane (*i.e.*, in *cis,* Figure 4A) or between antigens in closely apposed membranes (*i.e.*, in *trans*). Although antigen crosslinking has been observed for some influenza-specific antibodies (16, 18, 37–41), it remains unclear how common this phenomenon is, and how it depends on the specific epitopes which an antibody binds.

**Figure 4.**
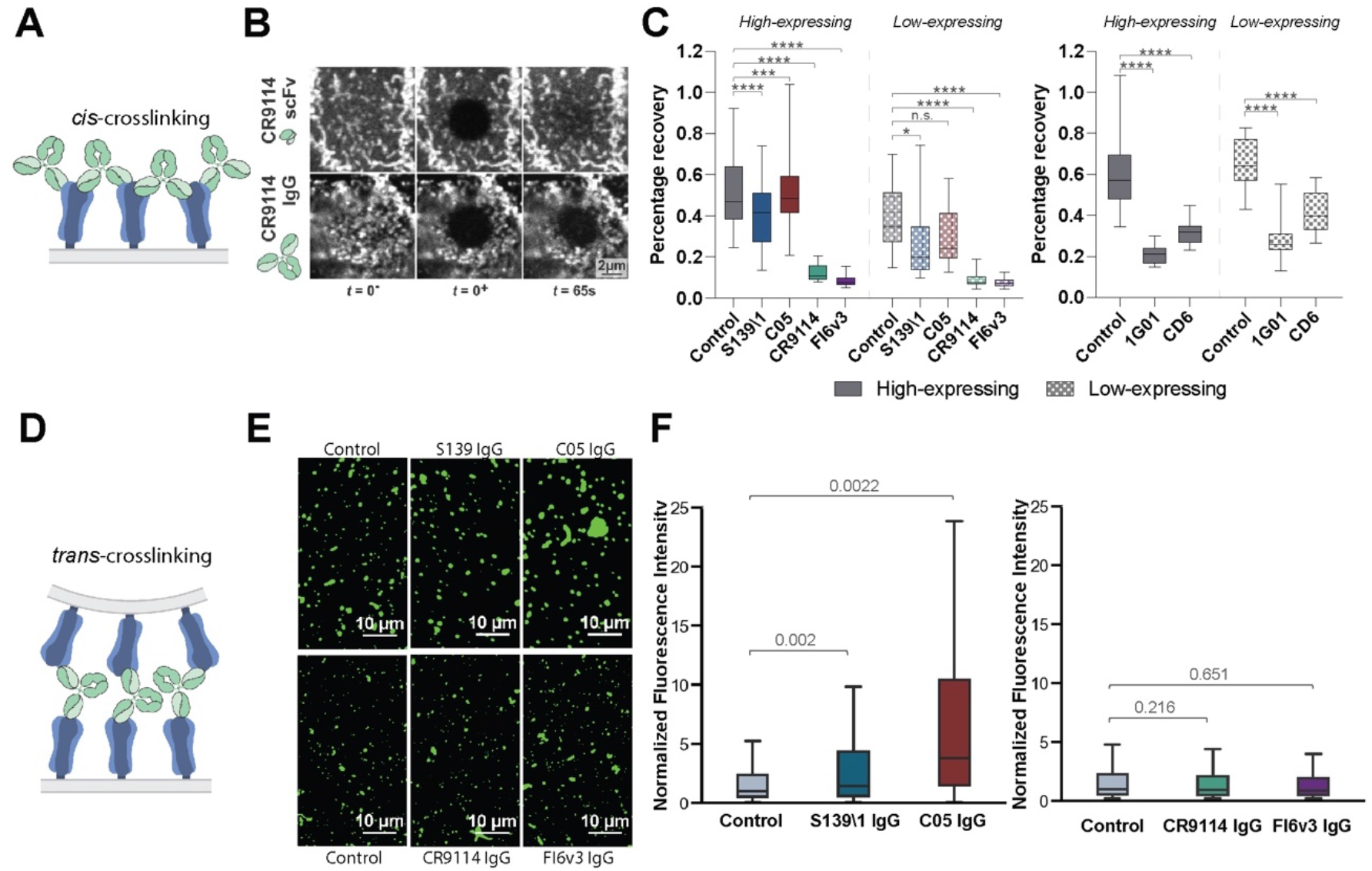
Antibodies inhibit virus assembly and release through distinct mechanisms. (A) Cartoon illustration of *cis* crosslinking of hemagglutinin (HA) by bivalent IgG antibodies. (B) Sample images of the apical cell surface immediately before and after photobleaching (*t* = 0^-^ and 0^+^) and at 65 s afterwards. (C, left) Percentage recovery of photobleached HA at 65 seconds post-photobleaching. Antibodies are tested at their respective IC75_release_ values. Data is combined from at least 50 photobleached cells per condition. Analyzed cells are split into high- and low-expressing groups based on the intensity of HA staining on the cell surface. (C, right) Percentage recovery of photobleached neuraminidase (NA) at 65 seconds post photobleaching. Data is combined from 30 photobleached cells per condition. *P* values are determined by Mann-Whitney tests. (* indicates p<0.05, ** indicates p<0.01, *** indicates p<0.001, **** indicates p<0.0001, ns indicates not significant.) (D) Cartoon illustration of *trans* crosslinking of HA by bivalent antibodies. (E) Sample images of virus particles/aggregates. (F) Distributions of particle/aggregate size, measured via fluorescence intensity. Antibodies are tested at their respective IC75_release_ values. Fluorescence intensities are normalized to the median value of their respective control groups. Results from RBS-binding and stem-binding antibodies are plotted separately because different non-competing fluorescent Fabs are used to measure particle/aggregate size. Data is combined from 3 biological replicates using stocks of A/WSN/1933 expanded separately. *P values* are determined using the mean of individual biological replicates by paired t-tests.

*Cis* crosslinking of trimeric or tetrameric viral surface proteins by bivalent antibodies could result in extensive networks of proteins with reduced mobility, a scenario that can be readily detected using fluorescence recovery after photobleaching (FRAP) (42). To avoid changes in protein mobility that could arise in the context of productive infection, we performed FRAP on cells transfected with either HA or NA plasmids and treated with bivalent or monovalent targeting antibodies at ∼48 hours post transfection (Figure 4B). While HA bound by monovalent CR9114 scFv showed efficient recovery (60% after 75 s), HA bound by bivalent CR9114 or FI6v3 IgG at concentrations that inhibit 75% of viral release did not significantly recover (Figure 4C, left). In comparison, RBS-binding antibodies S139\1 and C05 IgG showed *cis* crosslinking only on cells expressing high levels of HA, with S139\1 having a stronger effect than C05 (Figure 4C, left). Similar experiments evaluating NA mobility in the presence or absence of 1G01 IgG or CD6 IgG demonstrate that both antibodies significantly reduce NA diffusion, consistent with *cis* crosslinking (Figure 4C, right).

We next investigated the ability of antibodies to crosslink HA across membranes in *trans* (Figure 4D). Previous studies have shown that RBS-specific antibodies can cause virus aggregation on infected cell surfaces or in suspension (16, 18, 37, 39, 40). To compare *trans* crosslinking across the antibodies in our panel, we incubated virus particles overnight with antibodies at concentrations that result in 75% inhibition of virus release and measured particle aggregation using fluorescence microscopy (Figure 4E). While C05 and S139\1 IgG resulted in significant aggregation relative to IgG-free controls, the stem-binding antibodies CR9114 and FI6v3 did not (Figure 4F). These observations suggest that membrane-distal epitopes support antigen crosslinking across membranes, while membrane-proximal epitopes restrict crosslinking to antigens within the same membrane.

### Antibodies have widely varying potencies in inhibiting viral entry and release

Many of the anti-HA antibodies from our panel have established functions in blocking viral entry, by preventing either attachment (S139\1, C05) or membrane fusion (CR9114, FI6v3, CR8020). We sought to determine how the potency of these antibodies at inhibiting viral entry (‘IC50_entry_’, measured using single-round microneutralization assays) compares to their potency at inhibiting viral release (‘IC50_release_’). We find that the ratio of IC50 values varies widely across antibodies (Figure 5A). For example, although S139\1 and C05 both inhibit viral attachment, the two antibodies differ ∼10-fold in their IC50 ratio: specifically, C05 is similarly potent in inhibiting viral entry and release while S139\1 is ∼10-fold more effective at blocking viral entry than viral release. When we tested these antibodies against HAs towards which they have different affinities (HK68 and WSN33), we found that the ratio between IC50_entry_ and IC50_release_ remained similar for both antibodies (Figure 5B, Table S1). Thus, while the absolute potency of an antibody at inhibiting viral release depends on its affinity, its relative potency at inhibiting entry versus release appears to depend on other factors. Interestingly, inhibition of virus shedding by S139\1 IgG decreases at antibody concentrations above ∼5 nM. This is consistent with a transition from bivalent to monovalent binding as antibodies in solution begin to compete with bound antibodies for free HAs, disrupting *cis* or *trans* crosslinking (43).This phenomenon may limit inhibition of viral release for antibodies with exceptional affinity.

**Figure 5.**
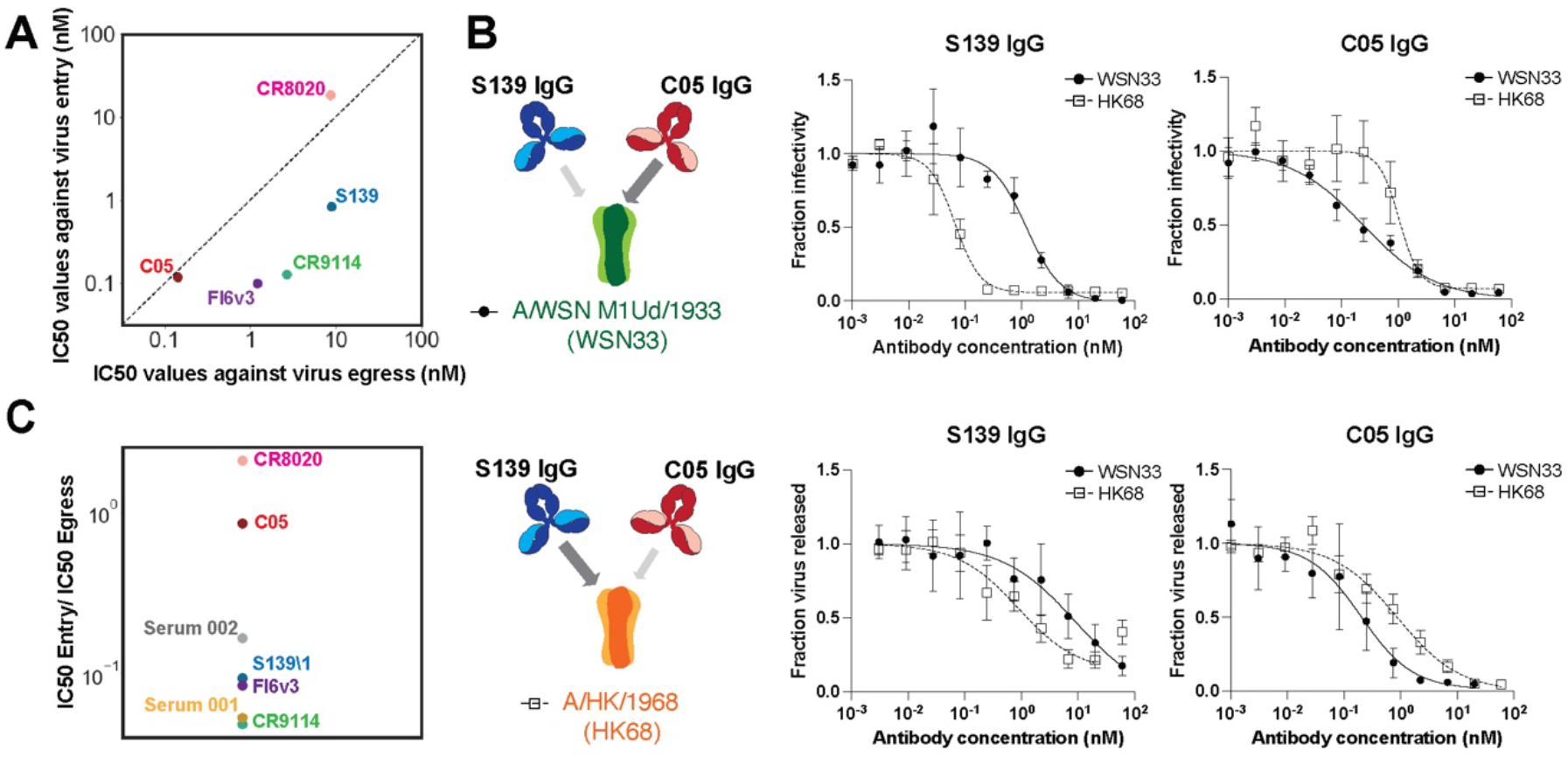
Antibodies and human sera differ widely in their ability to inhibit viral entry and viral egress. (A) Plot showing IC50 values for antibody inhibition of viral release and entry for the classically neutralizing antibodies from Figure 2. (B, Left) Illustration of binding preferences of S139\1 and C05 for different HAs. Wider arrows indicate stronger binding. (B, top right) Neutralization curves for S139\1 and C05 against viral entry. (B, bottom right) Neutralization curves for S139\1 and C05 against viral release. (C) Ratio of IC50_entry_ to IC50_release_ for antibodies and human convalescent serum. Data in panels *B* and *C* is from three biological replicates.

To investigate how results using monoclonal antibodies compare to polyclonal mixtures, we compared inhibition of entry and release by human convalescent serum against A/California/04/2009 at their respective IC50 values. Across two samples, we observe that one (Serum 1) is ∼20-fold more potent at inhibiting viral entry relative to viral release, while the other (Serum 2) is ∼6-fold more potent (Figure 5C). This is broadly consistent with our experiments with monoclonal antibodies and indicates that inhibition of viral entry does not necessarily predict potency in inhibiting viral release. Collectively, these data demonstrate that antibody inhibition of viral release plays a supporting role in limiting the spread of infection by a wide variety of antibodies, occasionally rivaling inhibition of viral entry even for antibodies that block receptor binding or membrane fusion.

### Antibody-HA structures predict a broad range of cis and trans crosslinking preferences

We reasoned that the binding orientation of an antibody could influence its geometric preference for crosslinking in *cis* or in *trans*. To investigate this possibility, we developed a geometric model using structures of Fab fragments bound to HAs of different subtypes. Using each structure to constrain the position for one of the antibody’s Fab arms, the model randomly samples potential configurations for the second Fab arm and evaluates the compatibility of each sampled configuration with *cis* or *trans* crosslinking (Methods, Figure 6A & B, Figure S7). This model focuses on the effects of antibody binding position and orientation without accounting for affinity, kinetics, or epitope accessibility – factors which are also likely important. Among the antibodies we tested, predictions from this model qualitatively agree with experimental measurements. C05 has a high propensity for *trans* crosslinking, while for the stem-binding antibodies CR9114, FI6v3, and CR8020, *cis* crosslinking is preferred (Figure 6B & C). Also consistent with our crosslinking data (Figure 4), S139\1 is predicted to have a reasonable propensity for both crosslinking modes. Among antibodies that we have not tested, F045-092 (44), L3A-44 (45), and CH65 (46) – antibodies which all bind in or around the RBS - are predicted to have a strong preference for *trans* crosslinking (Table S2). In contrast, antibodies predicted to have a strong preference for *cis* crosslinking bind to a range of antigenic sites, including adjacent to the RBS (S139\1 (47)) the central stalk (31.b.09 (48)), and the trimer interface (S8V2-37 (49)) (Figure 6C & D, Table S2). We selected one of these antibodies, 31.b.09, for further testing. Although 31.b.09 binds to an epitope that largely overlaps that of CR9114, the heavy and light chains are rotated ∼180° in the structures of these Fabs bound to the HA central stalk (Figure S8A), positioning 31.b.09 in a way that we reasoned would promote *cis* crosslinking and perhaps make it a relatively better inhibitor of virus assembly. While CR9114 inhibits viral entry ∼20-fold more potently than virus assembly and release, 31.b.09 shows the opposite trend, inhibiting assembly and release more effectively than viral entry (Figure S8B). Collectively, the general agreement between structure-based predictions and our experimental results suggest that antibody binding orientation constrains crosslinking propensity and provides a metric for predicting inhibition of viral assembly and release.

**Figure 6.**
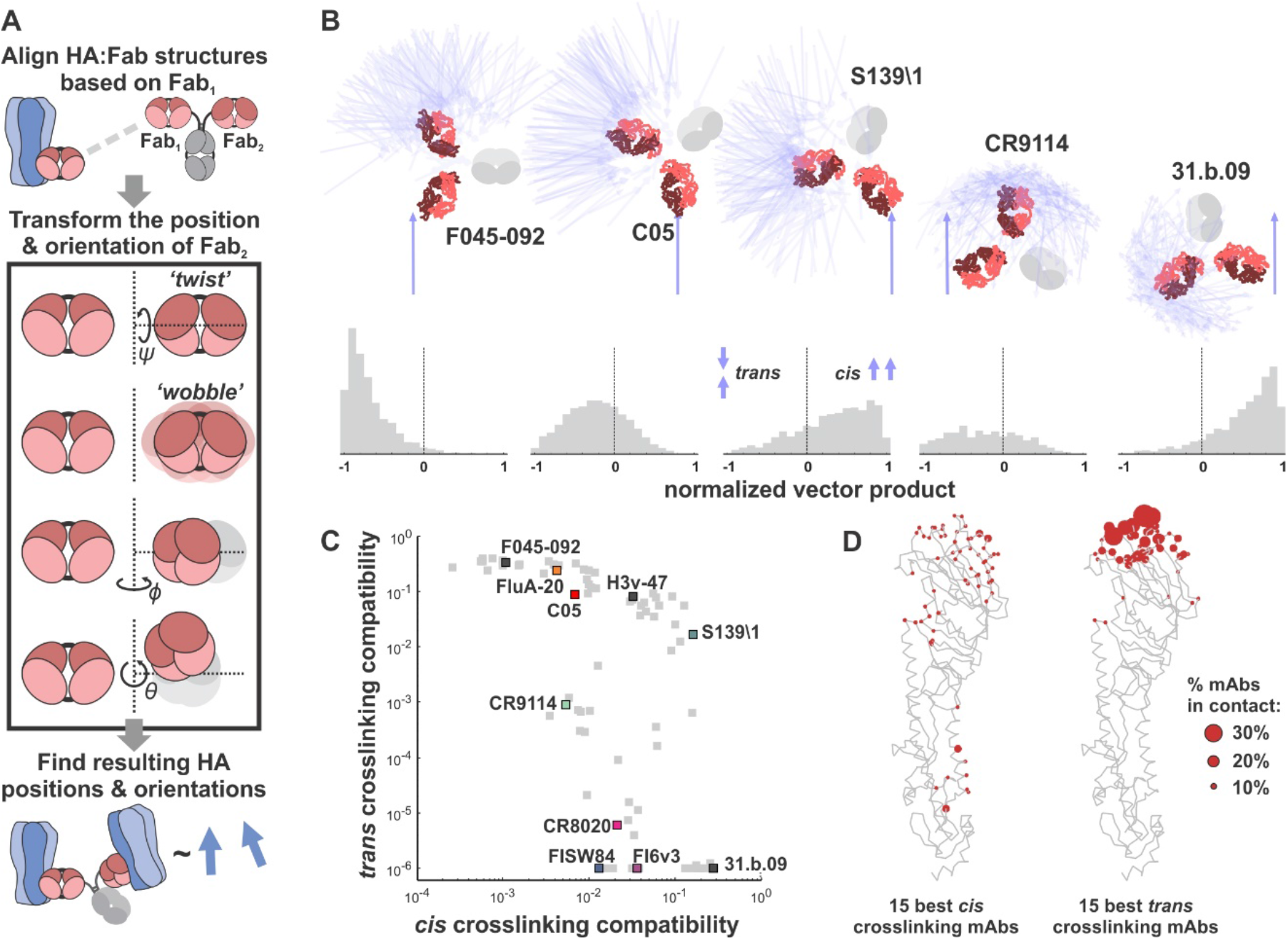
A structure-based model predicts *cis* and *trans* crosslinking for anti-HA antibodies. (A) Schematic overview of the modeling approach illustrating the transformations used to sample potential configurations of the two antibody Fab fragments (Fab_1_ and Fab_2_) and the resulting positions and orientations of the bound HAs. (B) Simulation results for selected antibodies. Top: Bound HAs are represented as blue vectors, Fab fragments are shown in red, and the antibody Fc (not included in the simulation) is shown in grey. Bottom: Distributions of relative HA orientations for the depicted antibodies. *Trans* crosslinking corresponds to anti-parallel HA orientation while *cis* crosslinking corresponds to parallel HA orientation. The resulting HA distributions are subsequently filtered to remove sterically forbidden configurations (Figure S7). (C) Predicted *cis* and *trans* crosslinking compatibilities for Fab:HA structures. Select antibodies characterized in this work or otherwise of note (F045-092, H3v-47) are highlighted. (D) Distribution of contact sites for the 15 best *cis* crosslinking (left) and *trans* crosslinking (right) antibodies based on the analysis shown in panel *C*. The backbone of an HA monomer is shown in gray. The size of the red spheres indicates the percentage of top antibodies that contact particular residues.

## DISCUSSION

While antibody inhibition of virus release has been documented in the context of infection by influenza and other enveloped viruses (50, 51), it remains an understudied aspect of antibody function. For the most part, antibodies that inhibit viral release have been identified via a process of elimination, through their inability to block viral attachment or entry despite restricting viral replication *in vitro*. By systematically and quantitatively comparing inhibition of virus release by a variety of neutralizing antibodies targeting distinct antigenic sites on HA, NA, and M2, we establish the generality of this phenomenon and identify distinct mechanisms through which antibodies restrict viral shedding: by crosslinking viral surface proteins in *cis,* to disrupt the diffusion of viral surface proteins; or by crosslinking *in trans,* to induce viral aggregation or sequestration on the cell surface. In addition to reducing the number of virus particles released during infection, we find that antibodies can also alter the size or morphology of released particles, potentially influencing viral replication in other ways. While we have focused specifically on IgG1 antibodies, other isotypes abundant in mucosal environments - especially IgA and IgM - will likely increase *trans*-crosslinking potency and may extend this capability to antibodies that recognize membrane-proximal epitopes. We propose that inhibition of virus assembly may serve as an additional metric to evaluate antibody potency alongside traditional neutralization measurements and assays aimed at evaluating Fc-dependent effector functions.

While the neutralizing activities of antibodies during viral entry are often well-correlated with their binding affinity, inhibition of assembly and release appears to depend on additional factors. One factor identified here is the antibody binding geometry. We find that the ability of antibodies to bridge two antigens - either in the same membrane (*cis*) or in opposing membranes (*trans*) – can be predicted with reasonable accuracy using a simplified computational model. This presents an opportunity to identify candidate antibodies that bind to neutralizing epitopes in favorable orientations that maximize crosslinking. We would expect that such antibodies would have dual potency in preventing viral entry as well as egress, given that their affinity is sufficiently high. A second factor that can contribute to inhibition of viral release is functional interactions between the target protein and other viral proteins; in particular, we find that the potency and mechanism with which anti-NA antibodies inhibit viral release depends on the HA and other viral proteins with which NA is paired. This result parallels early observations with anti-M2 antibodies, in which the potency of the ectodomain-specific antibody 14C2 was observed to differ markedly between strains despite complete conservation of the antibody epitope (9, 36). Understanding the polygenic nature of antibody inhibition of IAV release will require a deeper understanding of the mechanisms through which the IAV surface proteins contribute to assembly and budding.

We expect that many of the considerations that make an antibody a potent inhibitor of IAV assembly will also apply to other viruses that assemble at the plasma membrane of infected cells. Previous studies have reported that broadly neutralizing antibodies against E1 and E2 glycoproteins on alphaviruses are able to inhibit viral egress (52–56) (57). While crosslinking likely plays a role in this process, the ability of some E1-specific Fab fragments to inhibit alphavirus release similar to their IgG counterparts suggests that crosslinking is not a requirement (55). More generally, we speculate that viruses whose membrane proteins directly interact with each other on the viral surface or during assembly (58–60) may be vulnerable to inhibition by non-crosslinking antibodies that disrupt these interactions. Further investigation into the mechanisms through which antibodies restrict the assembly and release of influenza and other viruses may help guide the development of more broadly protective therapies and vaccines.

## MATERIALS AND METHODS

### Cell Lines and Viruses

MDCK-II and HEK-293T cell lines used in the study were purchased as authenticated cell lines (STR profiling) from ATCC and cultured with cell growth medium comprised of Dulbecco’s modified Eagle’s medium (DMEM; Gibco) supplemented with 10% fetal bovine serum (FBS; Gibco) and 1x antibiotic-antimycotic (Corning) under standard conditions (37 °C and 5% CO_2_).

Recombinant viruses were rescued using standard reverse genetics techniques (61). In brief, co-cultures of HEK-293T and MDCK-II were transfected with plasmids containing each vRNA segment flanked by bidirectional promoters. Viruses were collected from co-cultures at around two days post transfection and plaque purified. Viral plaques were passaged at an MOI of ∼0.001 in MDCK-II cells in virus growth medium comprised of Opti-MEM (Gibco), 2.5 mg/ml bovine serum albumin (Sigma-Aldrich), 1 μg/ml L-(tosylamido-2-phenyl ethyl) chloromethyl ketone (TPCK)-treated trypsin (Thermo Scientific Pierce), and 1x antibiotic-antimycotic (Corning). The viral stocks were further expanded by passaging at low MOI.

To obtain A/WSN/1933 viruses with filamentous phenotype, the M1 sequence within the WSN M segment is replaced by that of A/Udorn/1972. Rescue and characterization of these viruses has been described previously (62). To obtain A/California/2009 reassortant viruses, HA and/or NA segments from A/California/2009 are transfected in combination with corresponding genetic segments from A/WSN/1933 or A/Puerto Rico/8/1934.

### Antibody Purification and Labeling

Sequences for the variable regions of antibody heavy and light chains were obtained from deposited antibody structures on the PDB and cloned into expression vectors to make full-length human IgG1 antibodies. Sequences for the heavy chain were modified with a C-terminal ybbR tag for enzymatic labeling (63) and, in the case of Fab fragments, a His6 tag for affinity purification using Ni-NTA Agarose Beads (Thermo Scientific Pierce). Full-length antibodies were purified using protein A agarose beads (Thermo Scientific Pierce). Antibodies were expressed in HEK-293T following transfection with heavy and light chains at >70% confluency. Cells were subsequently cultured for seven days in Opti-MEM with 1x Anti-Anti and with or without 2% FBS for Fab and IgG antibodies, respectively. Supernatants from the HEK-293T cells were collected for affinity purification. Full details on antibody purification and enzymatic labeling are described elsewhere (63).

Human convalescent sera was obtained through BEI Resources (NR-18964 and NR-18965 for Serum 001 and Serum 002, respectively) and used without further purification in virus counting assays and microneutralization assays. IC50 values for serum neutralization were measured as fold-dilutions from the initial undiluted stock.

### Virus Counting Assay

MDCK-II cells were seeded into 96 well plates as a monolayer. Cells were washed twice with PBS and infected at MOI ∼ 1 for two hours at 37 °C. At 2hpi, the inoculum was removed, and cells were washed vigorously with PBS twice. Antibodies were serially diluted in virus growth media and added to cells for an additional 6 hours at 37 °C. Unless otherwise indicated, media was supplemented with 0.1 U/ml CpNA (Roche) to minimize steric inhibition of NA by stem binding antibodies. At 8 hpi, the supernatant was collected, diluted to the appropriate viral concentration to assure linear quantification (Figure 1B), and mixed with enzymatically-labelled viruses with sulfo-Cy5 site-specifically conjugated to HA (prepared as previously described (62)). Sulfo-Cy5 labeled virus served as a loading control for more consistent quantification. Addition of either CR9114 scFv 488 (A/WSN/1933) or FI6v3 scFv 488 (A/Hongkong/1968 and A/California/2009 reassortant strains) allowed discrimination between sample virus and the enzymatically-labeled loading control. Samples were imaged with a Nikon Ti2 confocal microscopy system using a 40x, 1.3-NA objective. Images containing virus particles were then analyzed using spot detection in each channel to determine the total number of virions immobilized on the glass-bottom well. The ratio between the particle count in the experimental and control samples were calculated for each antibody concentration and normalized to the antibody-free condition to generate the neutralization curve.

Imaging plates for virus quantification were prepared by coating coverslip-bottom wells (Cellvis) with 0.18 mg/ml BSA-biotin in PBS overnight at 4 °C. The imaging plate was then washed with PBS twice and incubated with 25 μg/ml streptavidin (Invitrogen) in PBS for 2 h. The imaging plate was then washed twice with PBS and incubated with 25 μg/ml biotinylated Erythrina cristagalli lectin (Vector Laboratories) at room temperature for 2 hours. Finally, the imaging plate was washed twice with NTC buffer (100 mM NaCl, 5 mM CaCl_2_; 20 mM Tris pH 7.4) prior to adding virus samples.

As a comparison to the virus counting assay, we performed quantitative Western blotting using the same serially diluted input virus stock used to generate the assay validation curve (Figure 1B). The same volume of virus samples was used as input for the imaging assay and for Western blotting. The blot was incubated with a polyclonal anti-HA primary antibody (Invitrogen) overnight at 4 °C, and probed with an IRDye 800CW goat anti-rabbit secondary (Licor) at room temperature for 1 hour prior to scanning on a Licor Odyssey imager.

### Microneutralization Assay

MDCK-II cells were seeded into 96 well plates as a monolayer. Monoclonal antibodies were serially diluted and mixed with viruses at 4°C for 2 hours. Cells were washed twice with PBS prior to adding the antibody-virus mixture and incubated at 37 °C. At two hours, the antibody-virus mixture was removed, and cells were washed twice with PBS and replenished with virus growth media containing 0.125 U/ml cpNA to prevent delayed primary infection from residual virus. The cells were incubated at 37 °C for 6 hours and imaged using CR9114 scFv 488 or FI6V3 scFv 488 to label HA on the surface of infected cells. The number of HA-positive cells were counted using the Spot Detection function in the Nikon Elements Analysis software and normalized to the antibody-free condition to generate the neutralization curve.

### Virus Aggregation Assay

Freshly expanded A/WSN/1933 viruses were treated with 0.125 units/mL CpNA for 2 hours at 4 °C. Viruses were then incubated overnight at 4 °C with antibodies at their respective IC75 values for inhibition of virus release (Figure 2). Virus-antibody complexes were then immobilized via ECL onto glass-bottom imaging chambers and labeled with fluorescent non-competing CR9114 scFv or C05 Fab and imaged using a 40X, 1.3NA objective. Images of virus-antibody complexes were then segmented using Nikon Elements Software and the HA intensity from the fluorescent non-competing Fab or scFv was quantified. Only viruses visualized with the same antibody fragments are directly compared in Figure 4.

### Fluorescence Recovery after Photobleaching Assay (FRAP)

An 8-well chambered cover glass (Cellvis) was incubated with 10μg/ml Human Plasma Fibronectin (EMD Millipore) at 4°C for 20 minutes. HEK-293T cells were transfected with HA- or NA-expressing plasmids and split into the imaging chambers. At ∼48 hours post transfection, fluorescently labeled antibodies are added to each well at the measured IC75 value (Figures 2 & 3) to perform FRAP. CR9114 scFv and CD6 Fab are used as negative controls to measure the normal diffusion of HA and NA in the absence of bivalent antibodies. An Olympus FluoView FV1200 laser scanning confocal microscope with a 60x, 1.35 NA objective was used for photobleaching and image acquisition. Photobleaching was performed over a circular region 1.98 μm in diameter using maximum laser power. One frame was taken before bleaching and one frame was taken 1 s after the bleaching event. Fourteen frames in total were collected at 5 s intervals following the photobleaching event.

To validate that the FRAP measurement is not complicated by rapid antibody turnover, dissociation kinetics of CR9114 and CD6 Fab fragments to A/WSN/1933 HA or A/CA/2009 NA were measured to verify that the time scale for dissociation far exceeds that of HA or NA diffusion. For these measurements, freshly expanded virus was incubated with CpNA for 30 minutes at 37 °C and immobilized onto a coverslip-based flow chamber via sequential layering of biotinylated BSA, streptavidin, and biotinylated ECL. Fluorescently labeled CR9114 Fab or CD6 Fab was introduced into the channel at ∼5 nM and allowed to bind for 30 minutes. The Fab was then washed away with PBS, and the fluorescent signal from remaining bound Fab fragments was measured at 5 s intervals for 75 seconds (the duration of the FRAP experiment). Results from this analysis are shown in Figure S9.

### Measuring Virus Particle Size

Images of shed virus particles collected with Nikon TI2 confocal microscopy system using a 60x, 1.40NA objective were segmented and quantified to determine the major and minor axis length and HA intensity measured by fluorescent non-competing antibodies as previously described (62).

### Modeling Antibody Crosslinking from Structural Information

To model the propensity of different mAbs to crosslink HA in *cis* or in *trans* based on their binding orientation, atomic coordinates for 83 structures of non-duplicate Fab fragments bound to HAs of different subtypes were downloaded from the Protein Data Bank and aligned by the Fab heavy and light chains. Using this first Fab arm (‘Fab_1_’) as a reference, a range of possible configurations for the alternate Fab arm (‘Fab_2_’) were sampled through sequential translation and rotation transformations, including introducing symmetry-breaking ‘wobble’ in Fab_2_, as observed for IgG antibodies (64–66) (Figure 6A). Based on previous estimates from electron microscopy, Fab_2_ conformations using *ψ* = +/-60°, *ϕ* = +/-30°, and *θ* = 60° +/-30° were sampled, where ‘+/-‘ indicates the standard deviations of the distribution each angle is sampled from. These simplified parameters result in a Fab arm which can twist extensively about its major axis (67), and which can rotate extensively along the other principal axes as well, consistent with previous observations (64–66).

Each sampled configuration of Fab_1_ and Fab_2_ results in a relative position and orientation for the bound HAs which was evaluated for its compatibility with *cis* or *trans* crosslinking. Configurations where the stem-to-stem or head-to-head distances are less than 5nm are eliminated, and head-to-stem distances less than 10 nm are eliminated as well (Figure S7A). For *trans* crosslinking, the head of the second (inverted) HA was required to be above the head of the first (upright) HA, while for *cis* crosslinking, the head of the second HA was required to be above the base of the first HA. For each allowable configuration, the dot product of axial vectors that extend from the bottom to top of the two HAs was evaluated. This produced a distribution of values ranging from -1 (for anti-parallel HAs) to +1 (for parallel HAs) which was used to score the geometric propensity of a given mAb to crosslink HAs in *cis* and in *trans*; antibodies whose distribution of vector products is clustered at +1 are highly compatible with *cis* crosslinking, while antibodies where the distribution clusters around -1 are highly compatible with *trans* crosslinking (Figure 6B). Changing the range of angular distributions sampled to determine possible configurations for bound HAs influences the predictions quantitatively but not qualitatively. Figure S7B shows how the predicted *cis* and *trans* crosslinking propensities for each mAb change under different implicit flexibilities; the dominant effect is to increase the ability of most antibodies to crosslink in *cis*, with little predicted effect on their ability to crosslink in *trans*.

### Statistics and Replicates

Statistical analysis was performed using GraphPad Prism 9, MATLAB, and Python. No statistical methods were applied to predetermine sample size. Statistical tests used are indicated in each respective figure legend. Box plots may omit outliers that are beyond the limit of the y-axes for clear visualization, but these are included in statistical analyses. Biological replicates are defined as cells separately infected/transfected/treated and assayed as indicated.

### Data Availability

Analyzed data is available through the manuscript. Raw images will be uploaded to Image Data Resource (https://idr.openmicroscopy.org/) upon publication. Analysis code and code used for Figure 6 and Figure S7 will be uploaded to GitHub.

## ACKNOWLEDGEMENTS

This work was supported by institutional funds from Washington University in St. Louis; National Institutes of Health grants R21 AI163985 and R01 AI171445; National Science Foundation CAREER Award 2238165; and Burroughs Wellcome Fund CASI Award 1013923. The following reagent was obtained through BEI Resources, NIAID, NIH: Human Convalescent Serum 001 to 2009 H1N1 Influenza A Virus, NR-18964 and Human Convalescent Serum 002 to 2009 H1N1 Influenza A Virus, NR-18965.

## AUTHOR CONTRIBUTIONS

Y.H. and M.D.V. designed research; Y.H., Z.G., and M.D.V. performed research; Y.H., Z.G., A.N.B., and M.D.V. contributed new reagents; Y.H., Z.G., S.S., and M.D.V. analyzed data; Y.H. and M.D.V. wrote the paper.

## COMPETERING INTEREST STATEMENT

The authors declare no competing interest.

---

**Figure S1.**
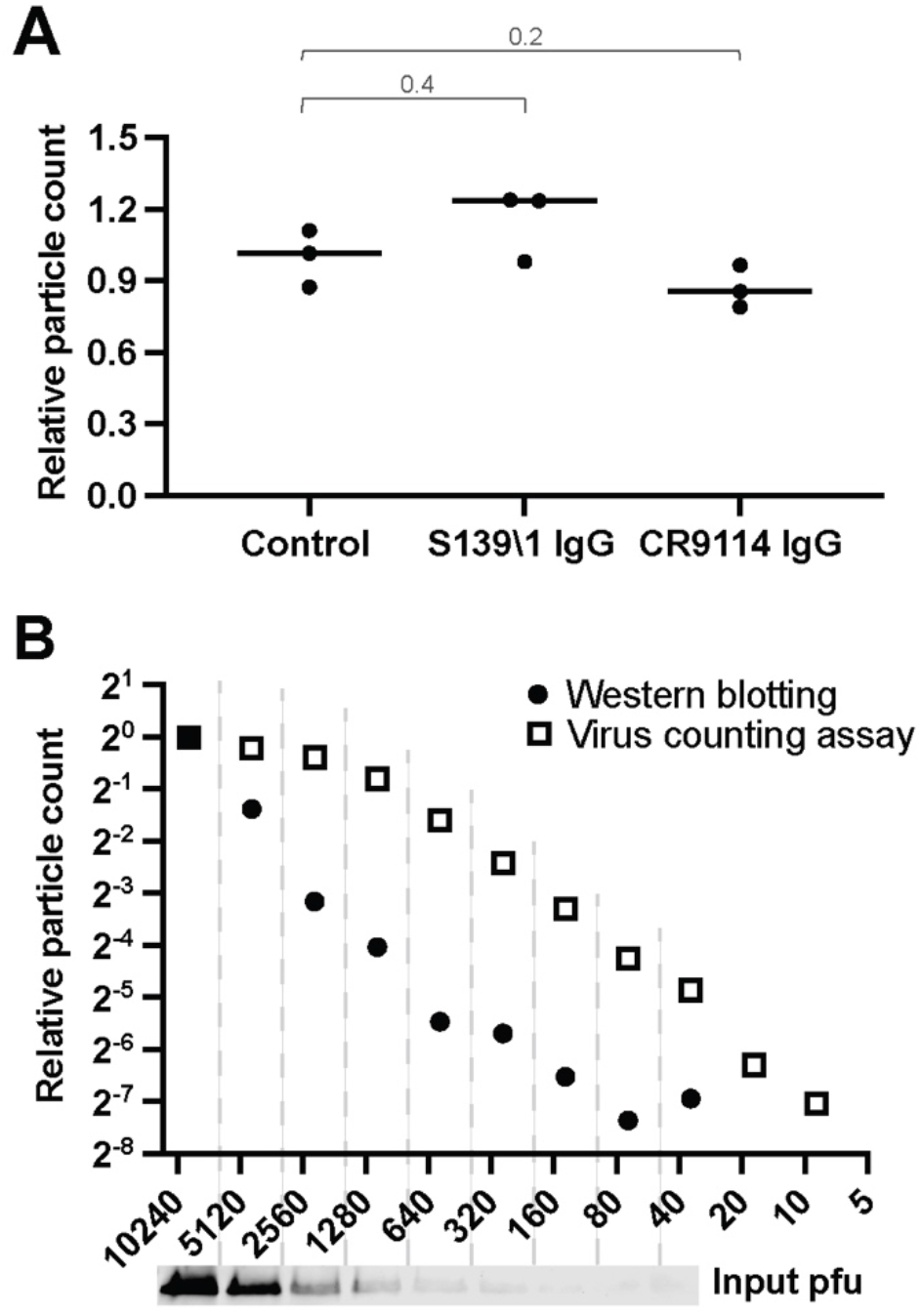
Evaluating robustness and sensitivity of image-based virus quantification. (A) A comparison of virus immobilization on ECL coverslips in the presence or absence of high concentrations of neutralizing antibodies. Antibodies are incubated with virus at a concentration of 60nM for 30 mins at 4°C before immobilization onto glass-bottom imaging chamber. Data is combined from 3 biological replicates and normalized to the mean of the control conditions. *P* values are determined by Mann-Whitney tests. (B) Comparison of particle counting results to Western blot analysis using an anti-HA antibody to quantify released virus. Results from western blot are collected from one serial dilution; results from the imaging-based assay are collected from three sets of serial dilution and the mean is shown. Contrast in the western blot scan is exaggerated to show the HA bands.

**Figure S2.**
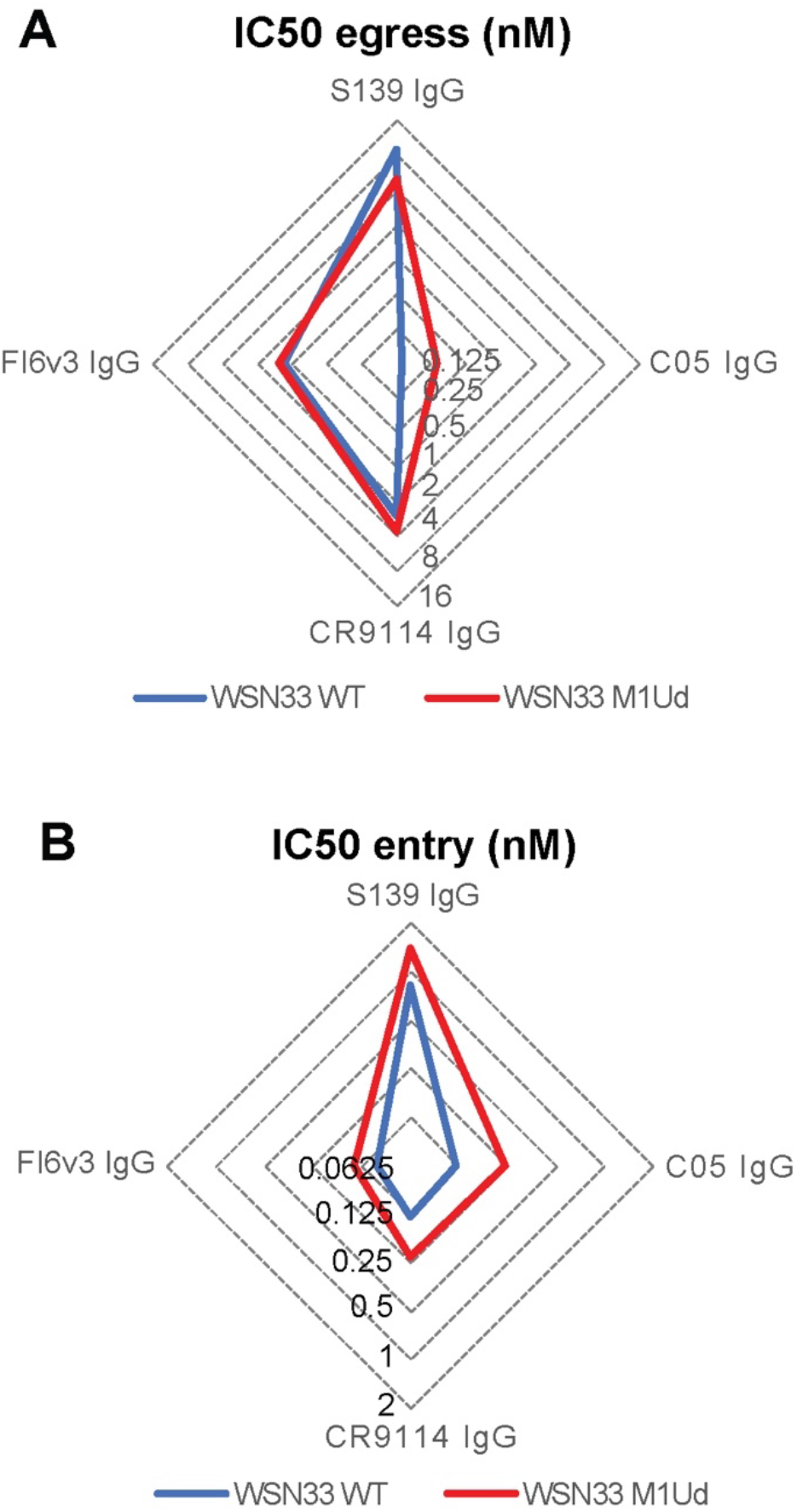
Inhibition of viral entry and egress differs between spherical and filamentous strains. (A) Radar plot showing the IC50 values for viral egress for spherical (WSN33 WT) and filamentous (WSN33 M1Ud) IAV strains with the same HA. (B) Radar plot showing IC50 values for viral entry (determined from microneutralization assays) for the same viral strains as in *A*. IC50 values for viral egress and viral entry are obtained from 3 biological replicates.

**Figure S3.**
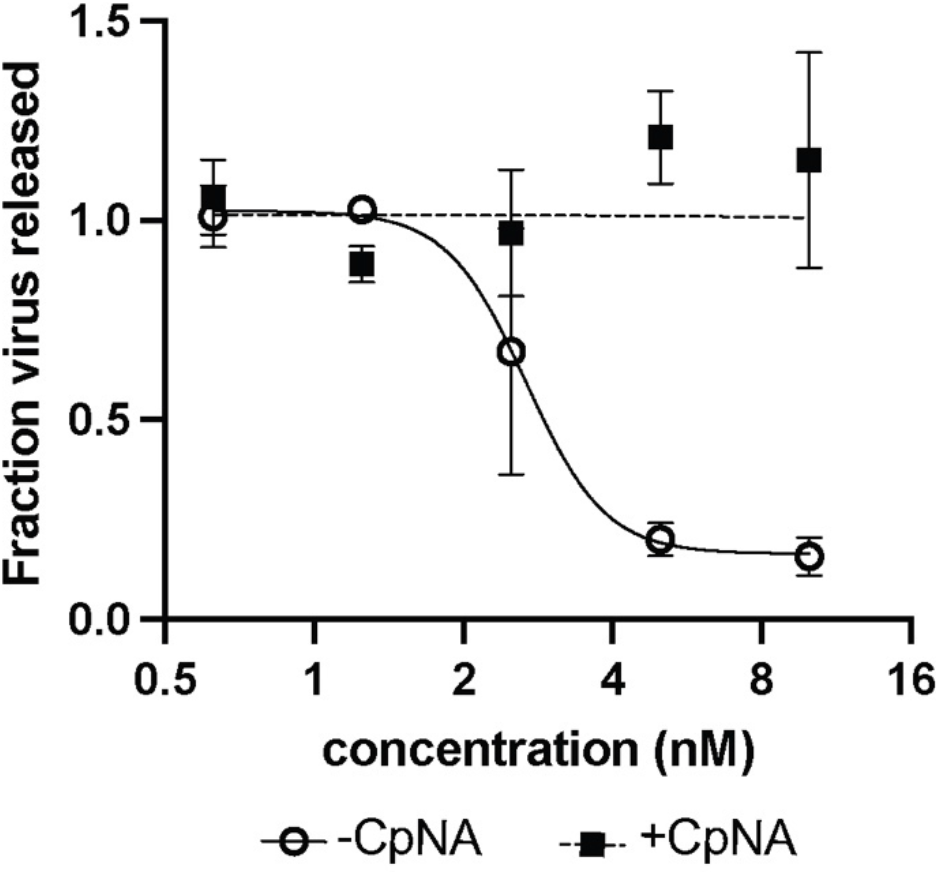
Exogeneous sialidase supplements NA activity. Inhibition of viral egress by oseltamivir carboxylate, with or without supplementation of exogeneous sialidase. Data is combined from 3 biological replicates. Dashed line is added as a guide to the eye. Error bars show standard deviations and the fit curve is obtained using the least squares method in GraphPad prism.

**Figure S4.**
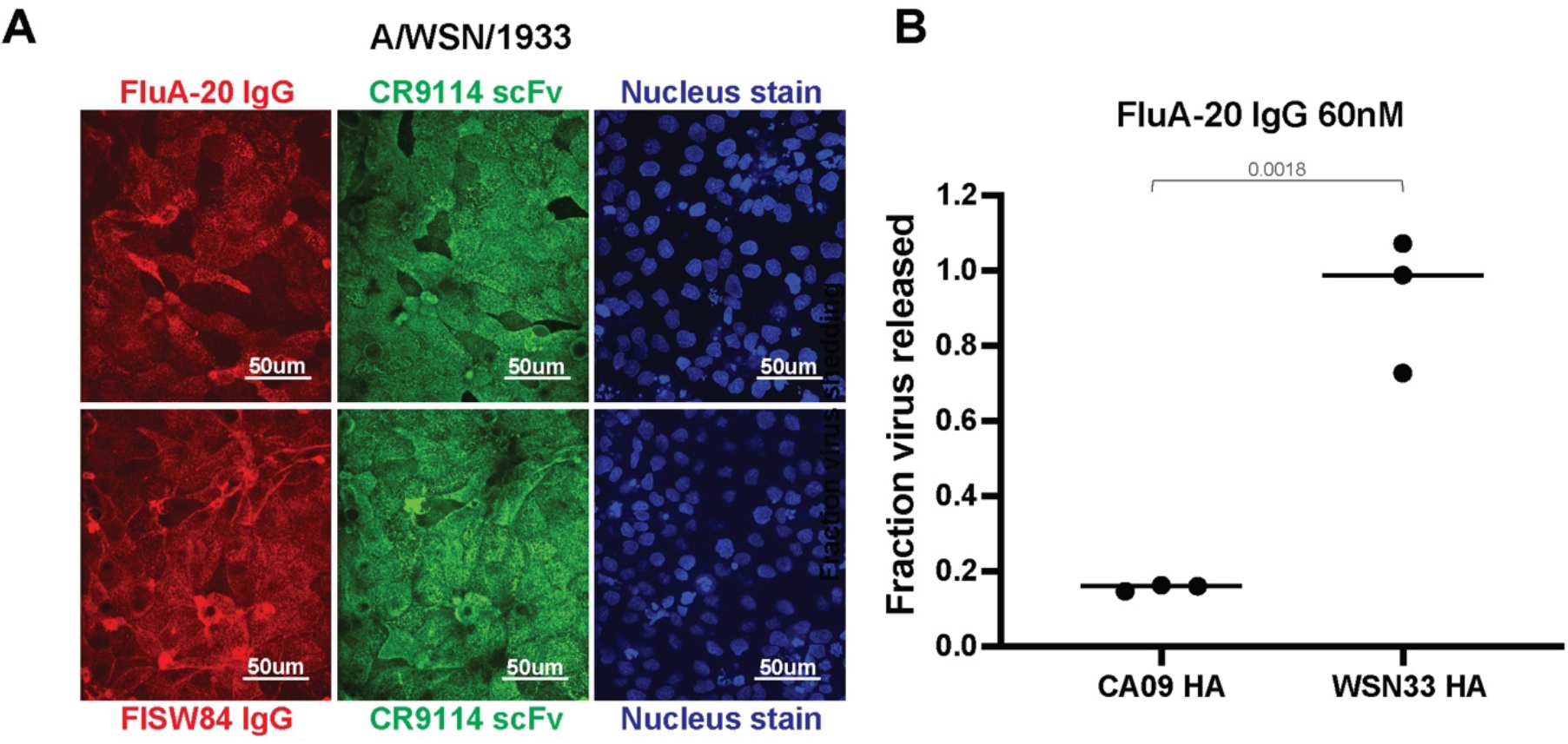
Binding and neutralization for antibodies with limited access to their epitopes. (A) Representative images of cells infected by A/WSN/1933 and treated with FluA-20 (top) or FISW84 (bottom) at 60nM for 6 hours post infection. Cell nuclei are stained for cell visualization.(B) Inhibition of viral shedding by FluA-20 for cells infected by IAV with HA from CA09 or WSN33. Data is normalized to the antibody-free condition. *P values* are determined by independent t-test.

**Figure S5.**
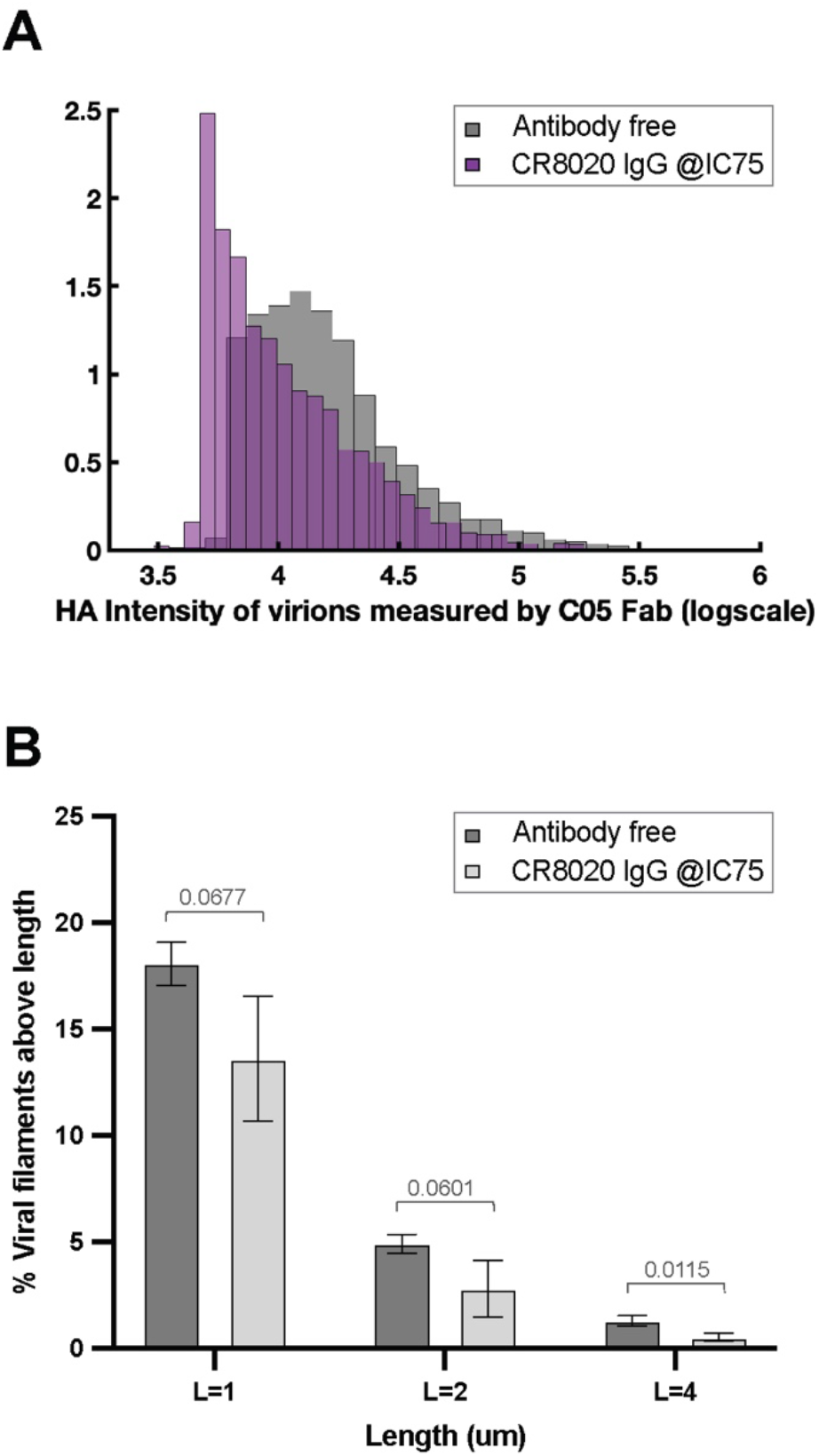
Effects of antibodies on the morphology of released virions. (A) Distribution of fluorescence intensities for individual viral particles shed in the presence or absence of CR8020 IgG. Fluorescence is measured using labeled C05 Fab. (B) Prevalence of viral filaments above different length thresholds released in the presence or absence of CR8020 IgG. Data is combined from 3 biological replicates. *P* values are determined by independent t-tests.

**Figure S6.**
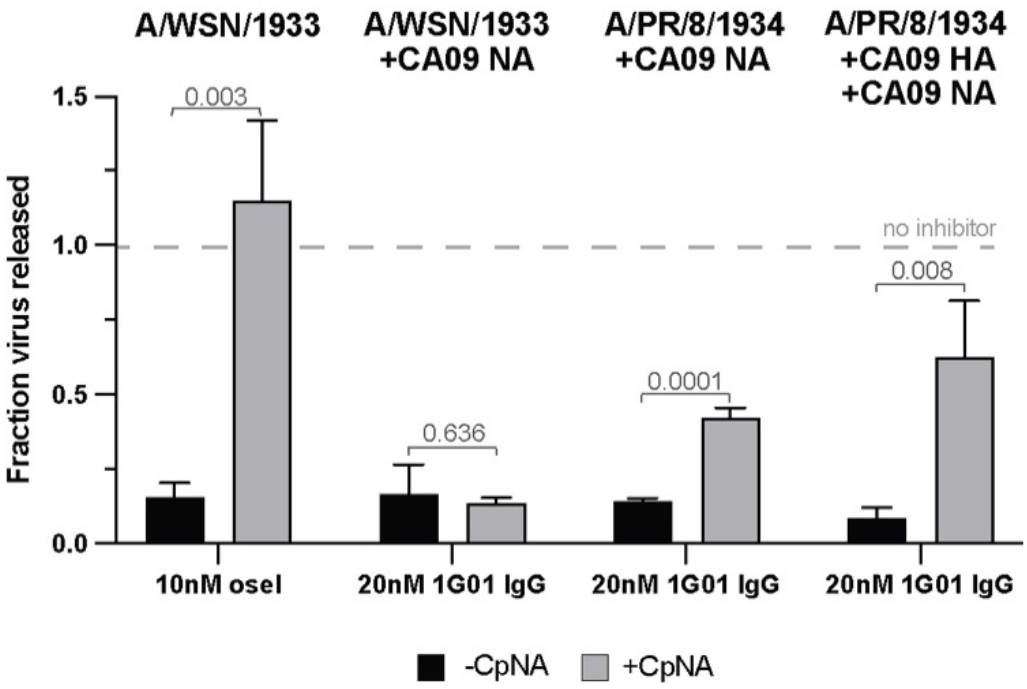
Inhibition of viral release by the anti-NA antibody 1G01 and oseltamivir. The plot shows the effect of NA inhibition by oseltamivir or 1G01 in the presence or absence of exogenous sialidase (CpNA) in different genetic backgrounds. Data for oseltamivir with or without CpNA, is repeated from Figure S3 using the A/WSN/1933 strain. Data is from 3 biological replicates. *P* values are determined by independent t-tests.

**Figure S7.**
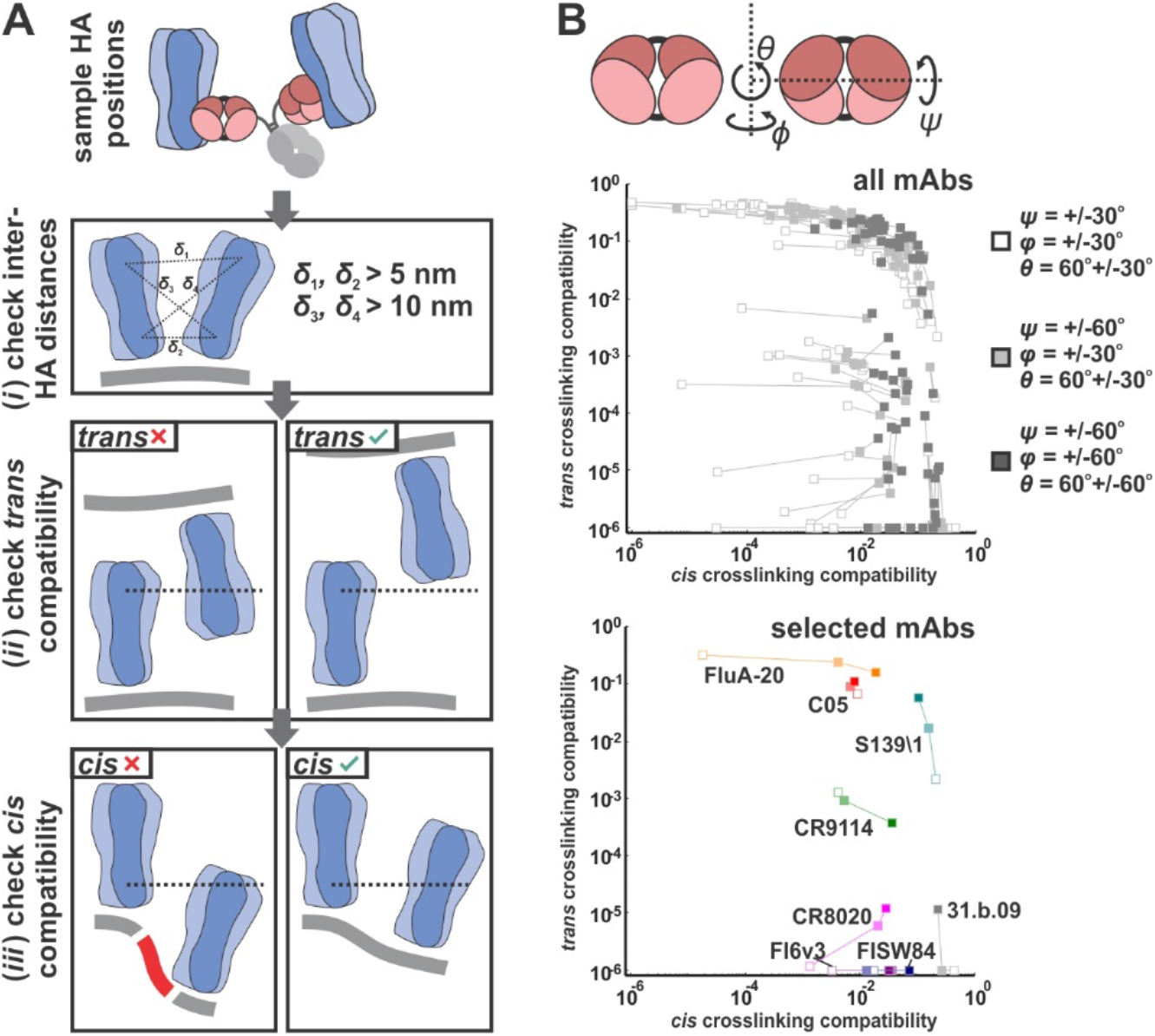
Evaluating *cis* and *trans* crosslinking with a structure-based model. (A) Schematic illustrating criteria for *cis* or *trans* crosslinking based on sampled positions and orientations of HAs. Candidate positions and orientations are sampled as shown in Figure 6A and described in Methods. (B) Predicted changes in *cis* and *trans* crosslinking propensities for antibodies modeled with different implicit flexibilities. Top: results for three models of flexibility across all 83 mAbs. Bottom: results for selected antibodies tested in this work.

**Figure S8.**
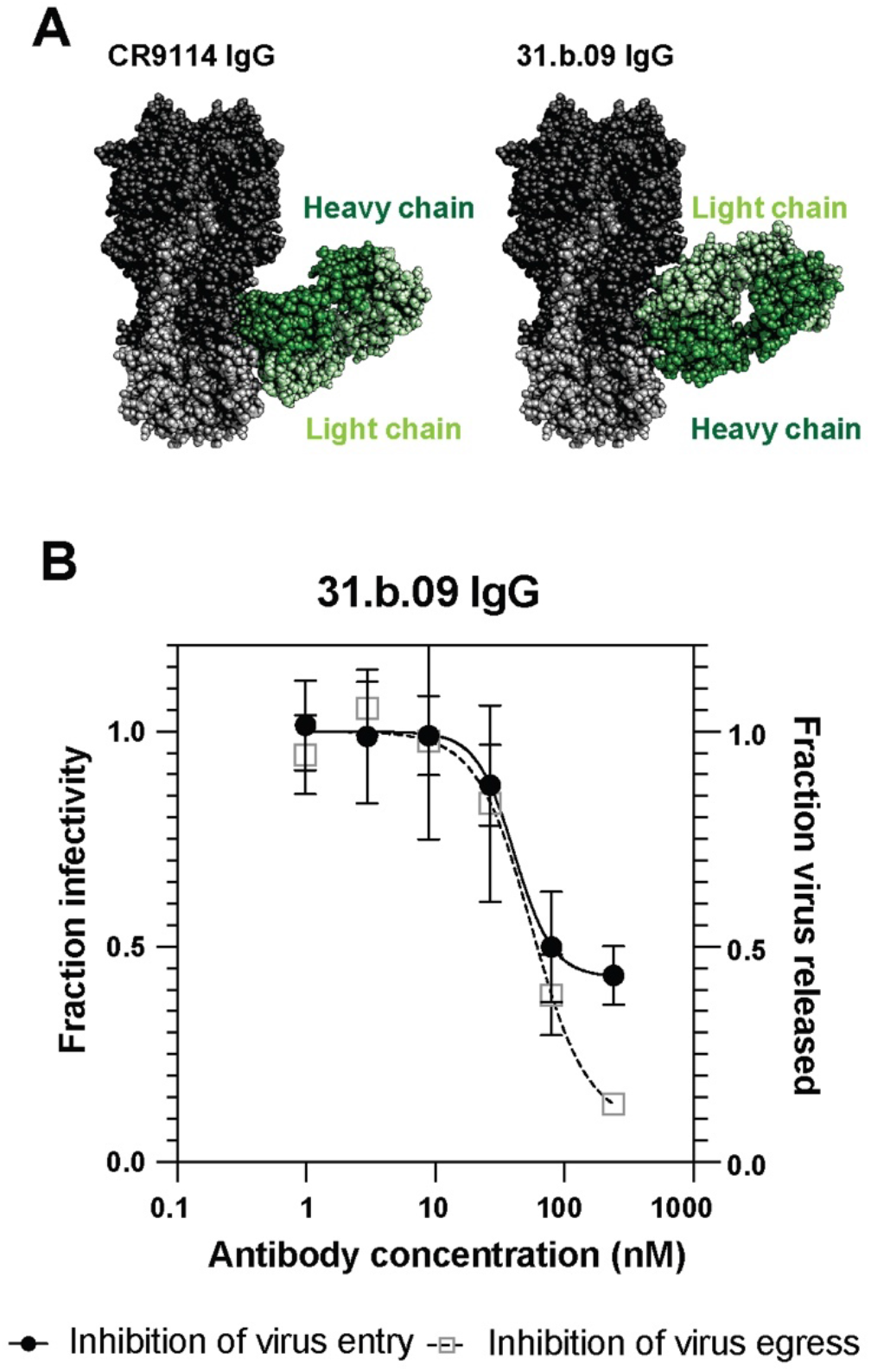
31.b.09 has similar inhibition potency against viral entry and egress. (A) Structure of CR9114 Fab (PDB ID 4FQI) and 31.b.09 Fab (PDB ID 5K9O) bound to HA. (B) Neutralization curves showing inhibition of viral entry and egress by 31.b.09 IgG against virus with CA09 HA and NA (PR8 reassortant). Curves are obtained from 3 biological replicates and normalized to the plateau of the fitted line for direct comparison. Error bars show standard deviations and the fit curve is obtained using the least squares method in GraphPad prism.

**Figure S9.**
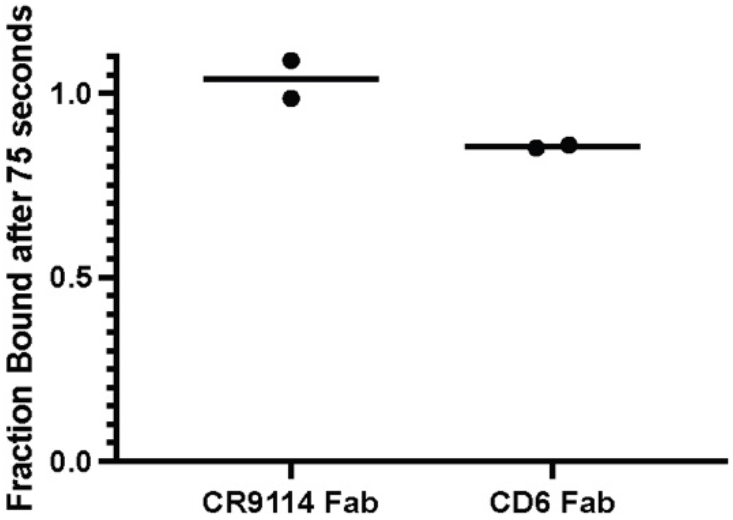
CR9114 Fab and CD6 Fab show minimal dissociation over the timescale of FRAP measurements. Measurement of CR9114 Fab and CD6 Fab dissociation from HA (WSN33) or NA (CA09) on the surface of virions. Data are from two biological replicates.

**Table S1.** Inhibition of viral entry and release for S139\1 and C05 against different viral strains. IC50_entry_ and IC50_release_ values against A/WSN/1933 (M1Ud) and A/HK/1968 are determined by least squares fit (GraphPad Prism) from data in Figure 5B. Both WSN33 M1Ud (labelled here as WSN33) and HK68 viral strains exhibit filamentous phenotype to facilitate comparison.

|  | HK68<br>entry<br>[nM] | HK68<br>release<br>[nM] | IC50 ratio<br>(HK68) | WSN33<br>entry<br>[nM] | WSN33<br>release<br>[nM] | IC50 ratio<br>(WSN33) |
| --- | --- | --- | --- | --- | --- | --- |
| S139\1 IgG | 0.066 | 0.868 | 0.076 | 1.221 | 8.961 | 0.136 |
| C05 IgG | 1.050 | 0.870 | 1.207 | 0.245 | 0.199 | 1.231 |

**Table S2.** Predicted cis and trans crosslinking propensities from HA-Fab structures. Data is plotted in Figure 6C and in Figure S7B. Scores less than 10^-6^ are set to 10^-6^ for visualization on a log scale. Please see *supplemental_table_2.xlsx* for complete data.

